# Highly Accelerated Vessel-Selective Arterial Spin Labelling Angiography using Sparsity and Smoothness Constraints

**DOI:** 10.1101/673475

**Authors:** Sophie Schauman, Mark Chiew, Thomas W. Okell

## Abstract

**Purpose:** To demonstrate that vessel-selectivity in arterial spin labeling angiography can be achieved without any scan time penalty or noticeable loss of image quality compared to conventional arterial spin labeling angiography.

**Methods:** Simulations on a numerical phantom were used to assess whether the increased sparsity of vessel-encoded angiograms compared to non-vessel-encoded angiograms alone can improve reconstruction results in a compressed sensing framework. Further simulations were performed to study whether the difference in relative sparsity between non-selective and vessel-selective dynamic angiograms were sufficient to achieve similar image quality at matched scan times in the presence of noise. Finally, data were acquired from 5 healthy volunteers to validate the technique in vivo. All data, both simulated and in vivo, were sampled in 2D using a golden angle radial trajectory and reconstructed by enforcing both image domain sparsity and temporal smoothness on the angiograms in a parallel imaging and compressed sensing framework.

**Results:** Relative sparsity was established as a primary factor governing the reconstruction fidelity. Using the proposed reconstruction scheme, differences between vessel-selective and non-selective angiography were negligible compared to the dominant factor of total scan time in both simulations and in vivo experiments at acceleration factors up to R = 34. The reconstruction quality was not heavily dependent on hand-tuning the parameters of the reconstruction.

**Conclusion:** The increase in relative sparsity of vessel-selective angiograms compared to non-selective angiograms can be leveraged to achieve higher acceleration without loss of image quality, resulting in the acquisition of vessel-selective information at no scan time cost.

## 1. Introduction

Angiographic methods are used to detect vascular abnormalities such as aneurysms, atherosclerosis, and arteriovenous malformations. However, many of the commonly used techniques for imaging the cerebral vasculature have associated risks. Digital subtraction angiography involves ionizing radiation and risk of complications because of the invasiveness of the procedure (1). In contrast enhanced magnetic resonance angiography (CE-MRA), there are also concerns associated with gadolinium-based contrast agents. Gadolinium-based contrast agents are unsuitable for patients with renal dysfunction because they can trigger nephrogenic systemic fibrosis (2), and have also been shown to be retained in the brain (3). Although the long-term health implications of gadolinium retention are still unknown (4,5), minimizing the use of gadolinium based contrast agents is recommended for clinical applications and research use.

Arterial spin labeling (ASL) can be used for non-contrast enhanced magnetic resonance angiography (NCE-MRA). Compared with other NCE-MRA methods, ASL is a more versatile and flexible technique. ASL can provide dynamic information about blood flow, as it is based on imaging a bolus of labelled blood just like in CE-MRA. Because of the complete removal of background tissue signal by subtraction of “label” and “control” images, smaller vessels can be resolved using ASL compared to, for example, time-of-flight angiography at the same resolution (6). ASL can also provide vessel-specific angiograms, whereas other magnetic resonance methods generally only provide non-selective angiograms. Vessel-specificity and dynamic information can be crucial in planning for surgical interventions and predicting clinical outcomes, for example in assessing collateral flow in stroke (7,8).

One way of achieving vessel-selectivity with ASL is using vessel-encoding (VE-ASL) (9), which is generally implemented as a variant of pseudo-continuous ASL (10). Compared to vessel-selective ASL methods that only label one vessel at a time (11,12), VE-ASL is more SNR efficient, as all acquired data informs all vessel-selective images. Hadamard encoding ensures that the SNR of a decoded VE image is the same as a for a non-vessel-encoded (nonVE) image acquired at the same scan time for fully sampled data (9). The main drawback of VE-ASL is longer acquisition times compared to nonVE-ASL because N+1 images are required to separate blood coming from N arteries compared with only a tag and control image for nonVE-ASL. To achieve matched scan time with nonVE acquisitions, the VE images have to be acquired with higher undersampling factors than the nonVE data.

However, we hypothesized that VE-ASL can be highly accelerated using non-linear reconstruction methods. Two main reasons why VE-ASL angiograms might be particularly well-suited to under-sampled reconstruction are related to intrinsic properties of angiographic data:

First of all, angiograms are spatially sparse. This can be exploited in a compressed sensing (CS (13,14)) acquisition and reconstruction framework. Compared with nonVE-ASL angiograms, VE-ASL images have higher relative sparsity because approximately the same number of non-zero voxels are distributed across multiple vessel-selective images. Because relative sparsity, along with image dimensionality and signal-to-noise ratio (SNR), contributes to the performance of a CS reconstruction (15) we hypothesized that VE-ASL angiograms could perform better than nonVE angiograms in a sparsity-constrained reconstruction.

Secondly, at sufficiently high temporal resolution the signal varies smoothly in time (16) as the bolus of labelled blood passes through the arterial tree. This temporal smoothness can be exploited to further regularize the underdetermined image reconstruction problem. While exploiting redundancy in the temporal domain is possible in the dynamic acquisitions provided by VE and nonVE ASL, non-dynamic methods like time-of-flight angiography cannot benefit from this extra dimension of information.

In this study, we present an accelerated acquisition and reconstruction method for VE-ASL angiography based on the enhanced spatial sparsity of vessel-specific angiograms and the smoothness of their temporal evolution. We demonstrate that the proposed method produces VE-ASL images of comparable quality to nonVE-ASL at matched scan duration at acceleration factors varying from R = 2 to R = 34, providing vessel-specific information at no additional cost.

## 2. Methods

### 2.1. Modelling the imaging system

The imaging system (Figure 1) was modelled as a linear equation:

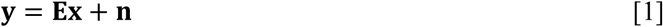

**Figure 1.**
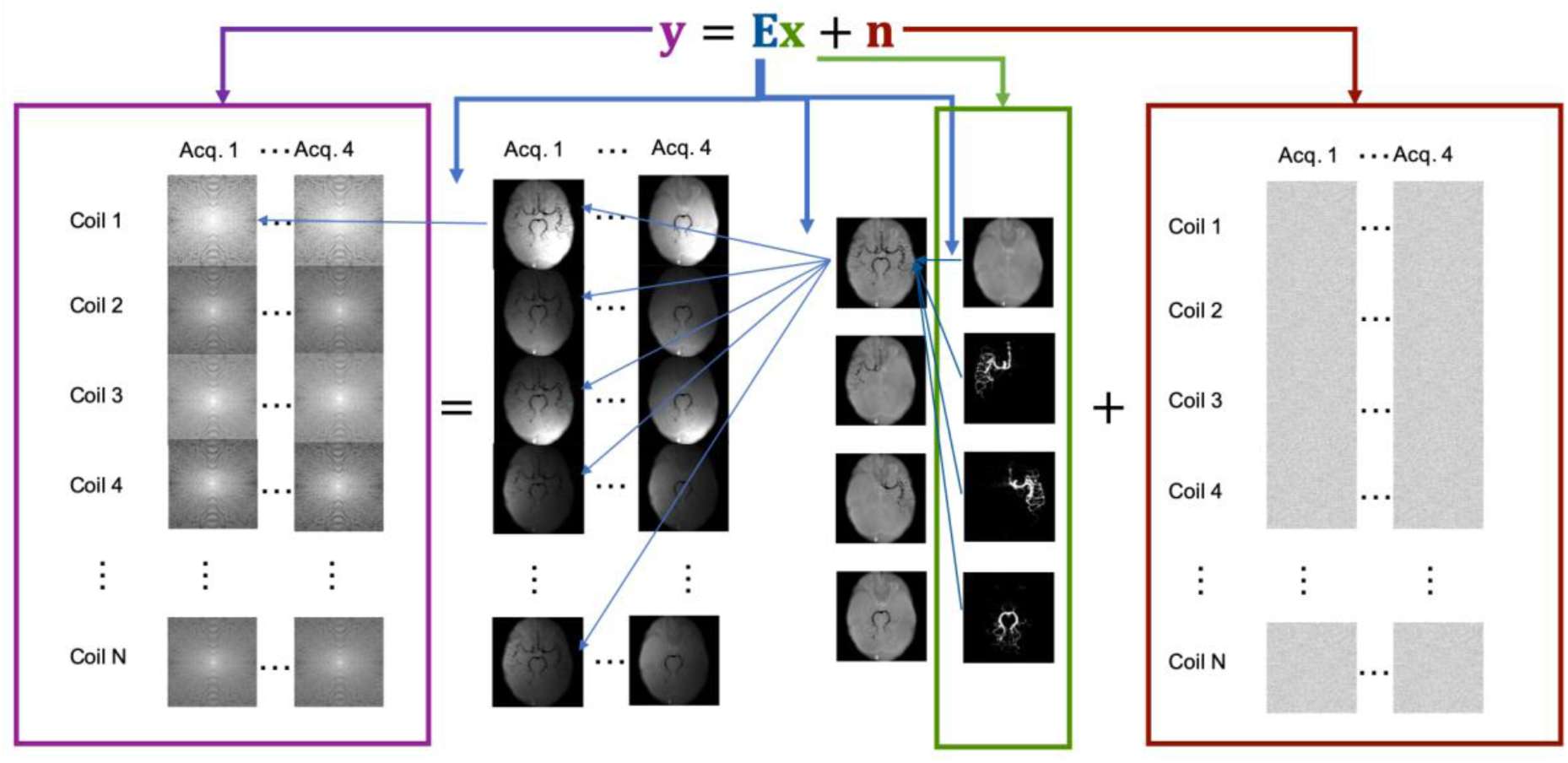
Model of imaging system used in simulations and reconstruction. y (purple box) represents the raw data that is a combination of noise, n (red box), and the object, x (green box), after it has been transformed by the imaging system, E (blue arrows), that consists of three parts (right to left): vessel-encoding, application of coil sensitivities, and an (under)sampled Fourier transform from physical space to k-space.

where **y** is a vector containing the complex signal measured by all the receive coils, with each entry representing one point in k-space in one coil. Noise, **n**, is a vector of complex white noise. The imaged object, **x**, is a vector containing the complex magnetization of blood from each vessel component as well as the static tissue for every position in physical space. Its length is therefore the number of voxels by the number of time points by the number of vessel components, i.e. a three-vessel VE image would have four components (three vessels and static tissue) and a nonVE image only two components (vessels and static tissue), thus making **x** twice as long in the VE case. In this work a three-vessel VE image was used, separating blood originating from three arteries, the right and left internal carotid arteries (RICA, LICA), and the basilar artery (BA). **E** is the linear operator that models the encoding of the magnetization. It contains three main parts: i) the linear combination of signal from blood and static tissue depending on the applied vessel-encoding scheme, ii) the multiplication of spatial sensitivity profiles for each of the receive coils in the system, and iii) the k-space sampling transform.

Simulations of the imaging system and subsequent reconstruction of both simulated and in vivo data was performed using MATLAB (Release 2017a, The MathWorks, Inc., Natick, Massachusetts, United States). The encoding operator (**E** in Eq. [1]) was implemented as a composition of functions instead of one single linear operator because of the large size of the problem. The vessel-encoding part of **E** was implemented directly as a matrix multiplication of a 2×2 Hadamard matrix (one image with labeled blood and one control image) for the nonVE case and a 4×4 Hadamard matrix for the VE case:

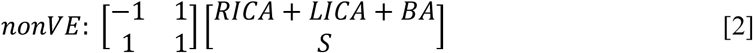

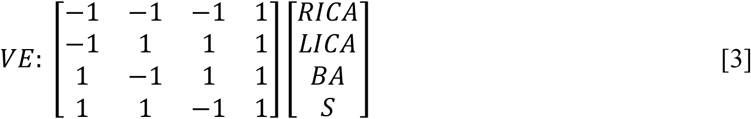

RICA, LICA, and BA represents signal from the blood coming from the respective arteries, and S represents the static tissue signal intensity.

The coil sensitivities and their conjugate transposes were applied as pointwise multiplication on the image or weighted combination of coils for the forward and adjoint transform respectively. The transforms between non-uniform k-space samples and image space were implemented using the non-uniform fast Fourier transform (NUFFT (17)) in the Michigan Image Reconstruction Toolbox (18). In our method we used a golden angle radial trajectory (19).

### 2.2. Simulations

Two datasets were used as ground truths for numerical simulations. One was a purely synthetic data set (numerical phantom) and the other a fully-sampled dynamic VE-ASL angiogram from a previous study (6). The numerical phantom consisted of a single frame of a hand-drawn ‘vessel-like’ image. It was used to confirm that the initial hypothesis (that increased relative sparsity improves the reconstruction) holds in a simplified system. The nonVE data had 14% non-zero values in the vessel image, whereas the VE data was only 5% non-zero in its three vessel images combined. In this simplified system the noise was set to zero and only one receive coil with uniform spatial sensitivity was modelled. The images were transformed into k-space data using the forward model described in section 2.1 and reconstructed with various amounts of undersampling (100%, 50%, 25%, 12.5%, 6.25% and 3.125% of the number of samples needed to reach the Nyquist limit with the golden angle radial sampling method (19)).

A real, high SNR angiogram was used to mimic the in vivo system as closely as possibly but with a well-defined ground truth and controlled noise conditions. Coil sensitivities previously measured using a phantom in a 32-channel head coil were included in the simulation. Complex Gaussian noise was added in k-space. k-Space SNR was defined as

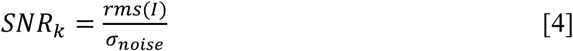

where rms(I) represents was the root-mean-square intensity of the noiseless k-space measurements and σ is the standard deviation of the added noise signal. Three noise conditions were simulated, i) no noise, ii) moderate noise (*SNR*_*k*_ = 185.7), and iii) high noise (*SNR*_*k*_ = 92.8). These simulated data sets were subsequently undersampled and reconstructed in the same manner as the in vivo data (see section 2.5 below).

### 2.3. In vivo acquisition

Five healthy volunteers (all male, age range: 25 to 34, mean age: 29) were scanned on a 3T Magnetom Verio (Siemens Healthineers, Erlangen, Germany) MRI scanner using a 32-channel head coil, with a dynamic 2D golden angle radial VE-ASL and nonVE-ASL sequence, similar to the implementation described in (20). All in vivo data was acquired under a technical development protocol approved by the local ethics committee.

Labelling was performed with bipolar pseudo-continuous ASL (9). The labelling plane was set just below the confluence of left and right vertebral arteries (VAs). For this study, the VAs were treated as a single artery to allow a four cycle Hadamard encoding scheme to be performed. More distal labeling where the VAs merge to form the BA was not performed to ensure artifact associated with the labeling plane did not overlap with the imaging region.

A spoiled gradient echo 2D golden angle radial readout was initiated (TR = 11.73 ms, TE = 5.95 ms, flip angle = 7°) immediately after the pCASL labelling pulse train (labelling duration: 1000 ms). For each preparation 108 radial spokes were read out during the 1266.8 ms long imaging period.

The same set of spokes were read out for each encoding before moving on to the next set of 108 spokes. The sequence used is shown schematically in Figure 2. The images were reconstructed with 1.1 mm x 1.1 mm x 50.0 mm spatial resolution, and 105.57 ms temporal resolution. The matrix size was 192 x 192. Each ASL preparation was preceded by a pre-saturation module for background suppression. A total of 34 ASL preparations were needed for each encoding to fill k-space to the Nyquist limit.

**Figure 2.**
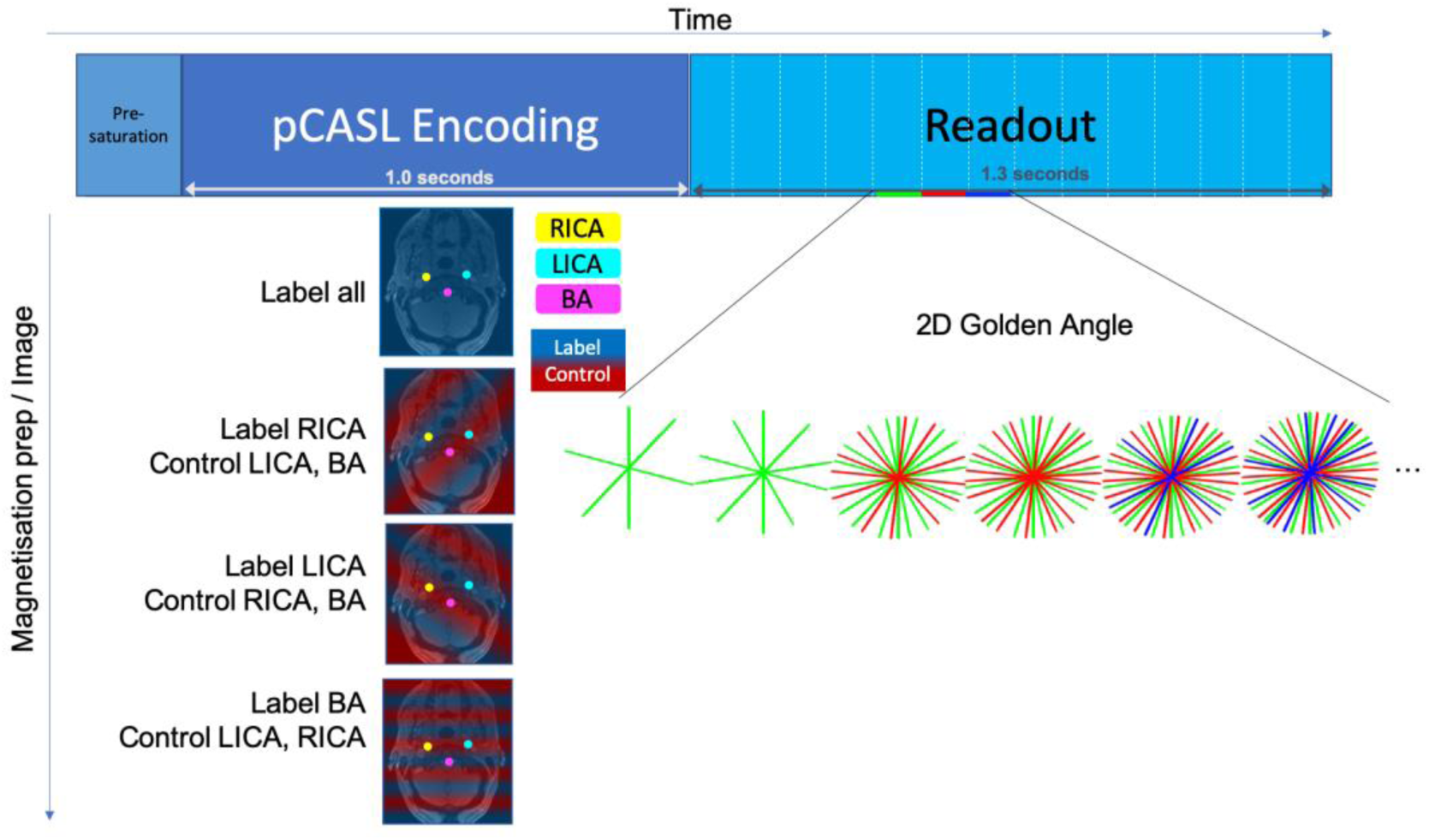
The imaging sequence consisted of a pre-saturation module for background suppression, pseudo-continuous ASL labelling and a spoiled gradient echo readout in a radial golden angle trajectory. The continuous readout was separated into frames in reconstruction. Four different magnetisation preparation modules were used to encode the left and right internal carotid arteries (RICA (yellow), LICA (cyan)) and the basilar artery (BA (magenta)). This colour scheme will be used in all subsequent figures. Transverse gradients within the pCASL pulse train generate spatial modulations to create tag (blue shading) and control (red shading) regions across the labelling plane.

For this study a 2D golden angle radial readout trajectory was chosen for a number of reasons. Firstly, for sparsity enforcing non-linear reconstructions it is important that the sampling pattern is as incoherent as possible to have noise-like aliasing artifacts (13). Radial imaging is more incoherent than Cartesian sampling and is thus better suited for heavy undersampling. The diffuse aliasing patterns of undersampled angiograms have also been exploited in similar work without compressed sensing or parallel imaging (21). Secondly, by changing the angle between the sampled spokes by the golden angle (approximately 111.25°) every additional spoke fills in the largest gap in k-space. This allows for flexibly choosing a subset of the data post acquisition by simply discarding a subset of the spokes, facilitating the evaluation of the reconstruction method at varying undersampling factors. It was also used to estimate the coil sensitivity profiles directly from the undersampled data.

The acceleration factor, R, was defined in relation to the fully Nyquist sampled acquisition time. An oversampled data set (acquisition time 10 min 53 s) for both the nonVE (R = 0.25) and VE (R = 0.5) cases were acquired to be used as ground truth. Then, independently acquired test data sets for both VE and nonVE (total acquisition time 5 min 26 s) were used to assess the reconstruction method. Before reconstructing, the test data were split into multiple subsets by grouping sequentially acquired spokes, such that the number of spokes in a group corresponded to a specific acceleration factor. The images were reconstructed at three different levels of acceleration with matched scan time between nonVE and VE: i) High undersampling - R=34 for VE (maximum acceleration as this only used one single ASL preparation for each encoding) and R = 17 for nonVE, scan time 10 s, ii) Medium undersampling - R = 8.5 for VE and 4.25 for nonVE, scan time 38 s, iii) Low undersampling - R = 2 for VE and R = 1 (no undersampling) for nonVE, scan time 2 min 43 s.

### 2.4. Pre-processing

To improve speed and reduce the memory burden of the reconstruction, the 32 coil channels were compressed to 8 channels using singular value decomposition (22). Because decoding of the signal was handled in the complex images, they are sensitive to phase errors. Therefore, for the in vivo data, phase correction was applied to account for B_0_ drift during the scan by minimising the average phase difference between the same spokes in different encodings with a scalar phase correction factor.

Coil sensitivity calibration images were reconstructed by combining k-space data across temporal frames to give one fully sampled or near fully sampled image. This was then used to estimate the relative coil sensitivity profiles for every point in space using the adaptive combine method (23). These estimated coil sensitivity profiles were used for generating the encoding operator, **E** as defined in Eq. [1], for each data set.

### 2.5. Reconstruction

Both the simulated and in vivo data were reconstructed using the same reconstruction method. Image reconstruction was achieved by the optimization of a non-linear cost function. Apart from data consistency, the cost function, *c*, included one term with the L_1_ norm of the (vessel-decoded) image to enforce image domain sparsity and one with the L_2_ norm of the finite difference along the temporal axis to enforce temporal smoothness:

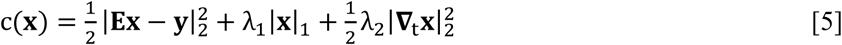

In the cost function **x** is the unknown image (the vessel components and the static tissue at all time points concatenated), **E** is the image acquisition operator, **∇**_t_ is the finite difference operator in the temporal domain, and **y** is the raw k-space data as defined in Eq. [1]. Here, the first term imposes data consistency of the reconstruction, the second term imposes sparsity, and the third term enforces temporal smoothness. λ_1_ and λ_2_ weigh the importance of the regularizing terms against data consistency. This cost function was minimized using the fast iterative shrinkage thresholding algorithm (FISTA (24)), using a Toeplitz embedding formulation to replace NUFFTs with FFTs for reduced computation time (25)

The appropriate regularization factors, λ_1_ and λ_2_ in Eq. [5], were determined experimentally by a grid search across a range of potential values. The (λ_1_, λ_2_) search space was chosen to be wide enough to ensure it fully characterized the target optima. The search space had to be varied for the different acceleration factors, as the absolute size of the data consistency term changed with how much data was acquired. λ_1_ was varied from 0 to 10^-5^ in steps of 10^-6^ for all acceleration factors both in vivo and for the simulated data. For the high undersampling (R = 17 and 34), λ_2_ was varied from 0 to 6 in vivo, and 0 to 2 in simulations, in steps of 0.2. For medium undersampling (R = 4.25 and 8.5), λ_2_ was varied 0 to 10 in steps of 1 for both the in vivo and simulation case. For low undersampling (R = 1 and 2) it was varied in steps of 2 from 0 to 20.

For the simulations, the combination of λ_1_ and λ_2_ with the highest correlation with the ground truth (as explained in section 2.6 below) for each acceleration factor and noise level was used. In vivo, the average performance across all subjects and eight subsets of data (except for the VE R = 2 and nonVE R = 1 case where only two subsets of data were acquired) were calculated and the regularization factors that produced the highest correlation score on average were chosen and used for all further analysis. Subject specific optimal regions (less than 1% quality reduction from the subject specific optimum) were also calculated to ensure that the chosen combination of regularizing factors was reasonable for all subjects.

### 2.6. Analysis

All reconstructions were compared against some ‘ground truth’ image. For the simulations, the actual input image was used directly for comparison. For the in vivo data the oversampled acquisition was reconstructed with minimal regularization applied for denoising (λ_1_ = 10^-6^, λ_2_ = 0) and used as ground truth.

For quantitative assessment of image quality, non-overlapping vessel-specific masks were applied to both the reconstruction and the ground truth as it was important that the quantitative assessment focused on the relevant pixels in the sparse images to avoid bias due to artefacts (e.g. from eye motion). The masks were generated by manually thresholding the temporal average image of the ground truth and then dilating that mask with a 3×3 structuring element with zeros in the corners and ones in every other position. The dilation ensured that both vessels and neighboring background was included into the mask. In voxels where multiple vessels provided signal (mixed blood supply), the voxel was assigned to the vessel with the dominant (highest intensity) signal. The masks were then applied to each frame of each vessel component (or the total vessel component for nonVE). The Pearson correlation coefficient (*r*) between the ground truth and reconstructed pixel values across all time points within the masks were then calculated. Qualitative assessment of reconstruction quality largely agreed with the metric.

When comparing nonVE against VE, the correlation coefficients, *r*, for each vessel mask were Fisher transformed to a *z*-score to make the distributions of correlation coefficients more Gaussian.

This then allowed Student’s t-tests to be performed to determine statistical significance at a 98.3% confidence interval (95% with added Bonferroni correction for multiple comparisons of the three vessel components).

## 3. Results

### 3.1. Effect of increased relative sparsity only in numerical phantom

In the fully simulated noiseless case on the numerical phantom, where the only difference between nonVE and VE was the ratio of non-zero to zero voxels, the VE data was reconstructed more robustly at higher undersampling factors than nonVE (Figure 3). When the data was fully sampled or near fully sampled, both VE and nonVE could be reconstructed essentially perfectly, but below 25% of Nyquist sampling, the reconstruction of the sparser VE simulation outperformed the nonVE version, achieving approximately matched performance for twice the undersampling factor, negating the factor of two time penalty that would be needed to perform three-vessel VE instead of nonVE angiography. The performance gain was also qualitatively visible in the reconstructed simulation images, where increased blurring and streaking artifacts were observed in the nonVE case.

**Figure 3.**
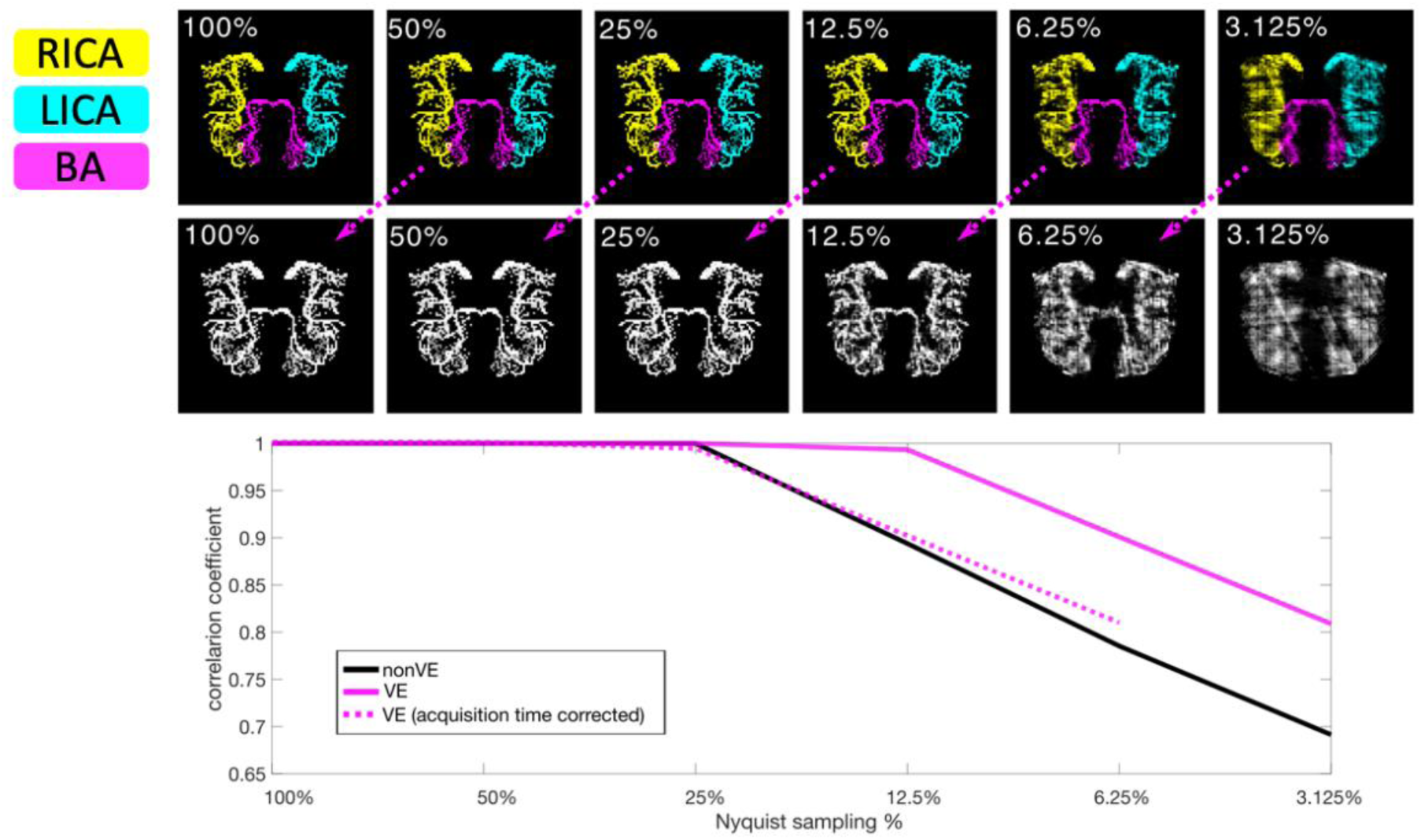
Numerical phantom simulations: Below 25% Nyquist sampling the quality of the nonVE reconstruction decreases rapidly. For VE this only occurs at twice the undersampling factor. The dashed line shows a shifted copy of the VE line to illustrate the reconstruction quality of a three-vessel VE angiogram at the same acquisition time as the nonVE. The corresponding time-matched images (top row) are linked via dashed arrows. Reconstruction quality is quantified using the correlation coefficient between voxels in the reconstructed image and the ground truth (100% sampling).

### 3.2. Simulations on real data

Similar results to the numerical phantom were observed in the real data simulations. With no added noise the VE and nonVE reconstruction quality was high (*r* > 0.99) at all acceleration factors, and no difference was found between VE and nonVE. With added noise and simulated matched scan times (equal SNR, but twice the undersampling for VE), VE was reconstructed marginally, but significantly, better (P < 0.01) than nonVE for low and medium acceleration factors in both medium and high noise conditions. At high acceleration factor, the results were more varied and VE performed better than nonVE in some vessels but worse in others and for some there was no statistically significant difference. A summary of the noisy simulation results is shown in Figure 4.

**Figure 4.**
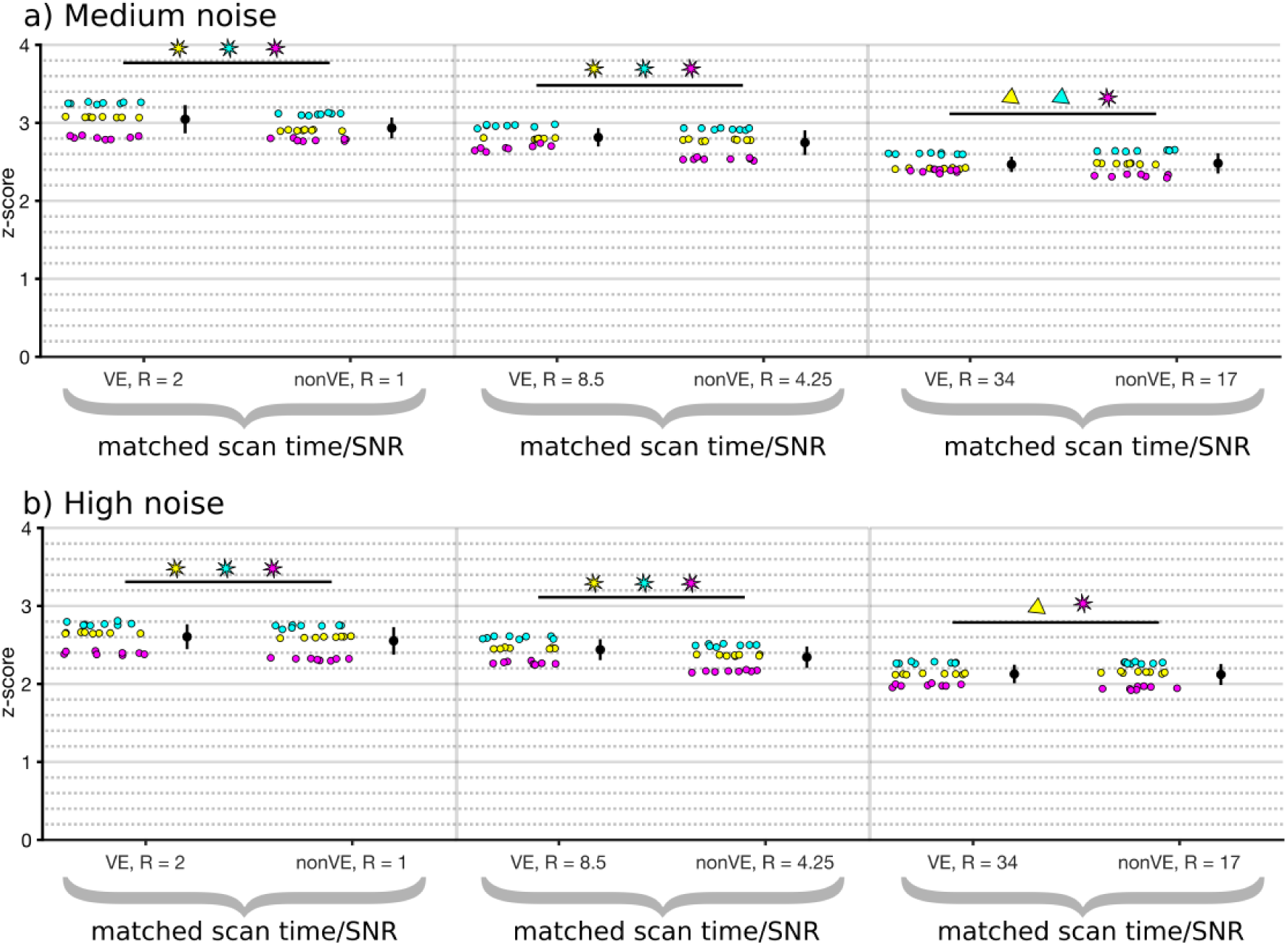
Reconstruction quality in simulations with a) medium noise, and b) high noise. Each scatter point represents the fisher transformed correlation coefficient calculated in a mask (RICA – Yellow, LICA – Cyan, BA – Magenta) for one reconstruction. Statistical significance between the time-matched nonVE and VE groups is represented by a star if VE had higher correlation coefficient and a triangle if nonVE did.

### 3.3. In vivo

#### 3.3.1. Optimal regularization factors

The optimal regularization factors for the in vivo reconstructions did not vary considerably between different subjects and their optimal ranges (within 1% from optimum) had considerable overlap at all acceleration factors (Figure 5).

**Figure 5.**
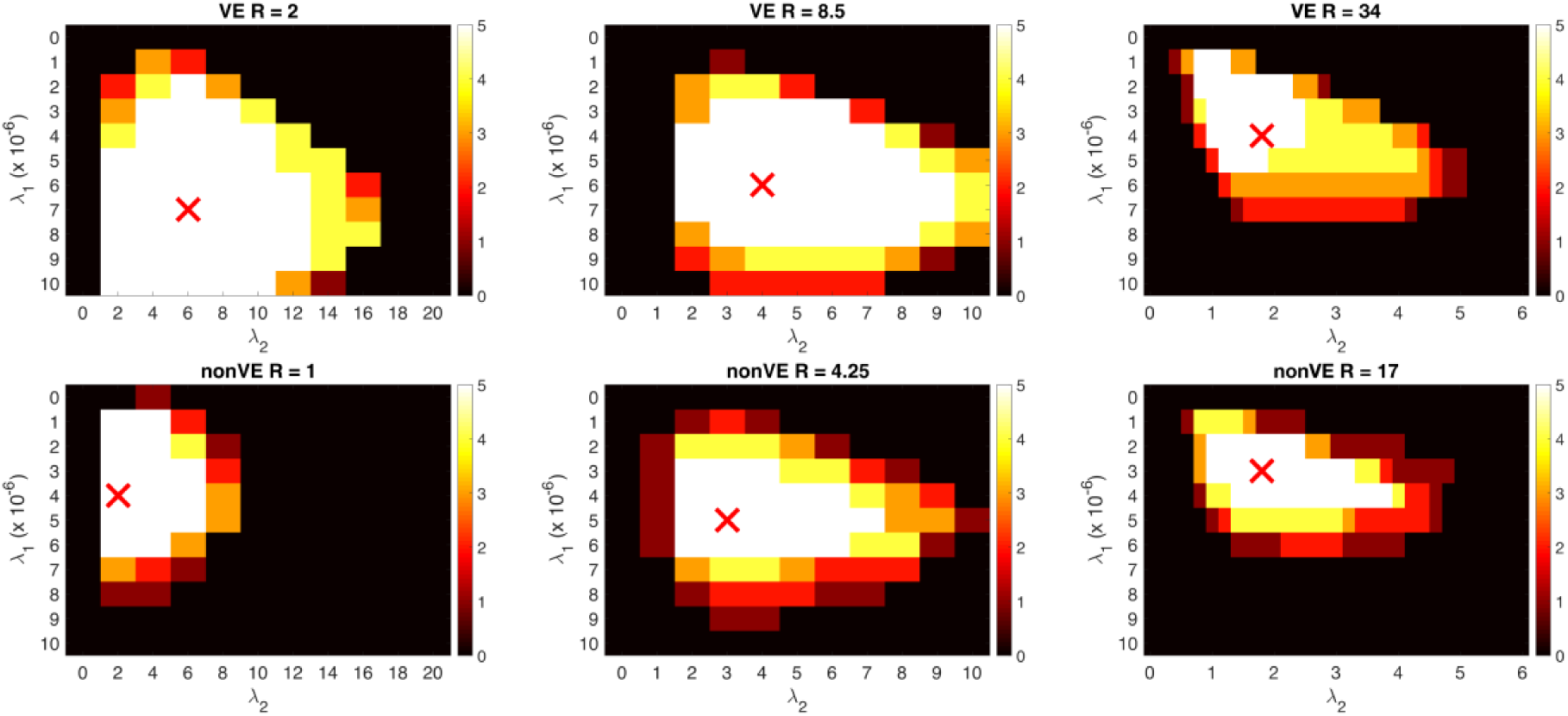
The optimal regularization factors (marked with red ‘x’) were within the optimal area (within 1% of optimum) for every subject. The color represents how many subjects had optimal reconstruction at each combination of regularization factors.

The effect of varying the regularization factor within the overall optimal range was varied sensitivity and specificity, i.e. reduced noise against the cost of losing visibility of small vessels. Some examples of this tradeoff are shown in Figure 6. For the sake of comparing VE with nonVE reconstructions in an un-biased way, the group optimum regularization factors for each acquisition method were used for all further analyses. The optimal regularization factors for each acceleration factor and subject/noise level are summarized in Supporting Information Table S1.

**Figure 6.**
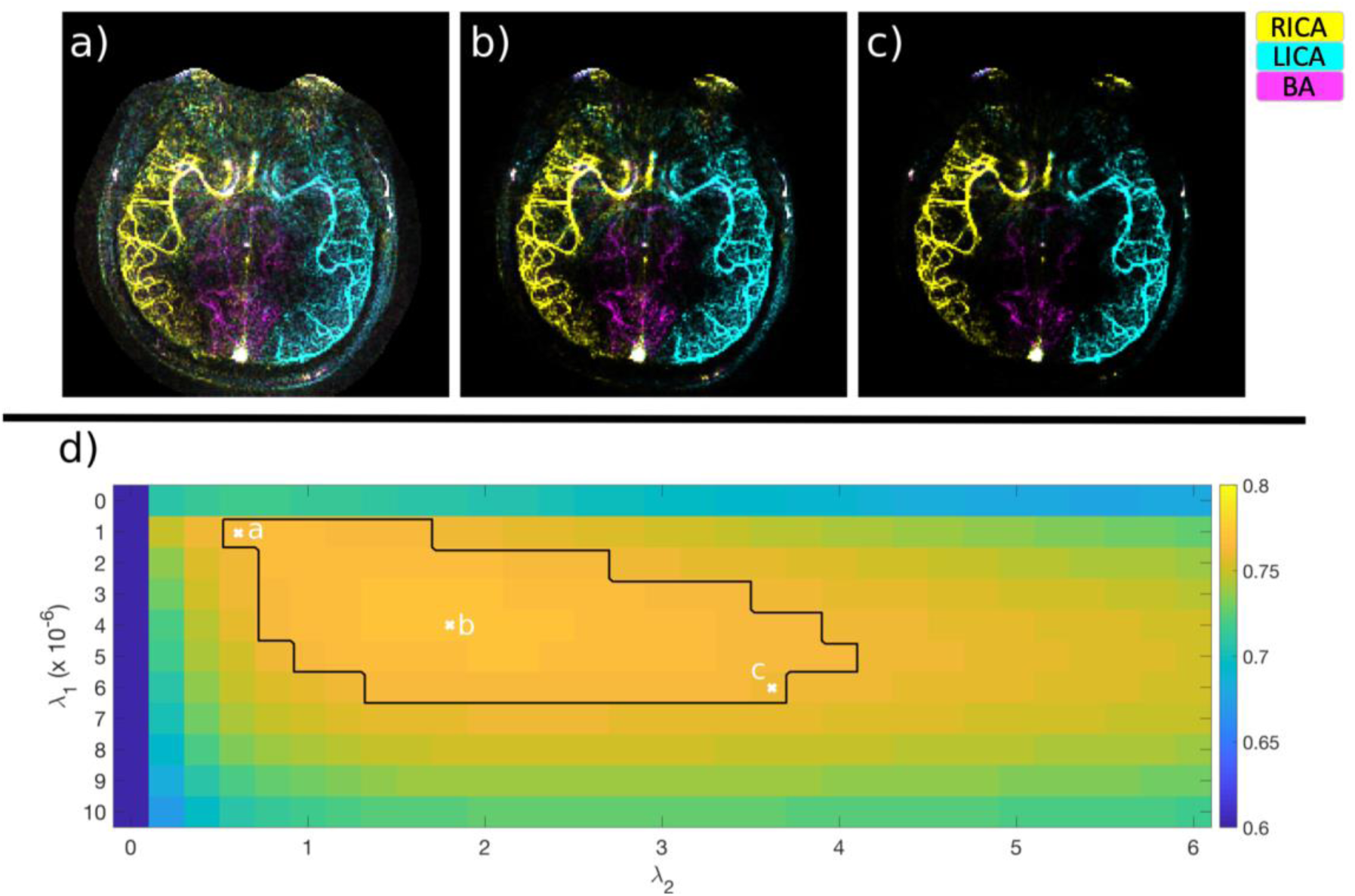
The same raw dataset (one of the in vivo VE R = 34 subsets) reconstructed with three different combinations of regularization factors, a) minimal regularization, more noise, b) optimal regularization based on the average correlation coefficient, c) maximal regularization, heavily denoised reconstruction. d) The average correlation coefficient across all subjects at each combination of regularization factors. The black border represents the area where the reconstruction was within 1% of optimum and the ‘x’s mark the regularization factors used for a), b), and c).

Both regularization terms improved the overall reconstruction quality in all cases. Average correlation coefficients for the in vivo data improved 4.6-55.2% by having a non-zero λ_2_, 2.3-8.9% by having a non-zero λ_1_, and 15.3-98.7% by having both regularization factors non-zero. Similarly, in simulations with non-zero noise a 1.0-13.0% improvement was observed for non-zero λ_2_, 0.1-10.0% improvement for non-zero λ_1_, and 2.2-88.3% improvement by having both regularization factors non-zero. Visually, the value of having both regularizing terms is clearly shown in an example reconstruction in Figure 7, with non-zero λ_2_ causing better delineation of the vessels, and non-zero λ_1_ reducing noise and noise-like artifacts.

**Figure 7.**
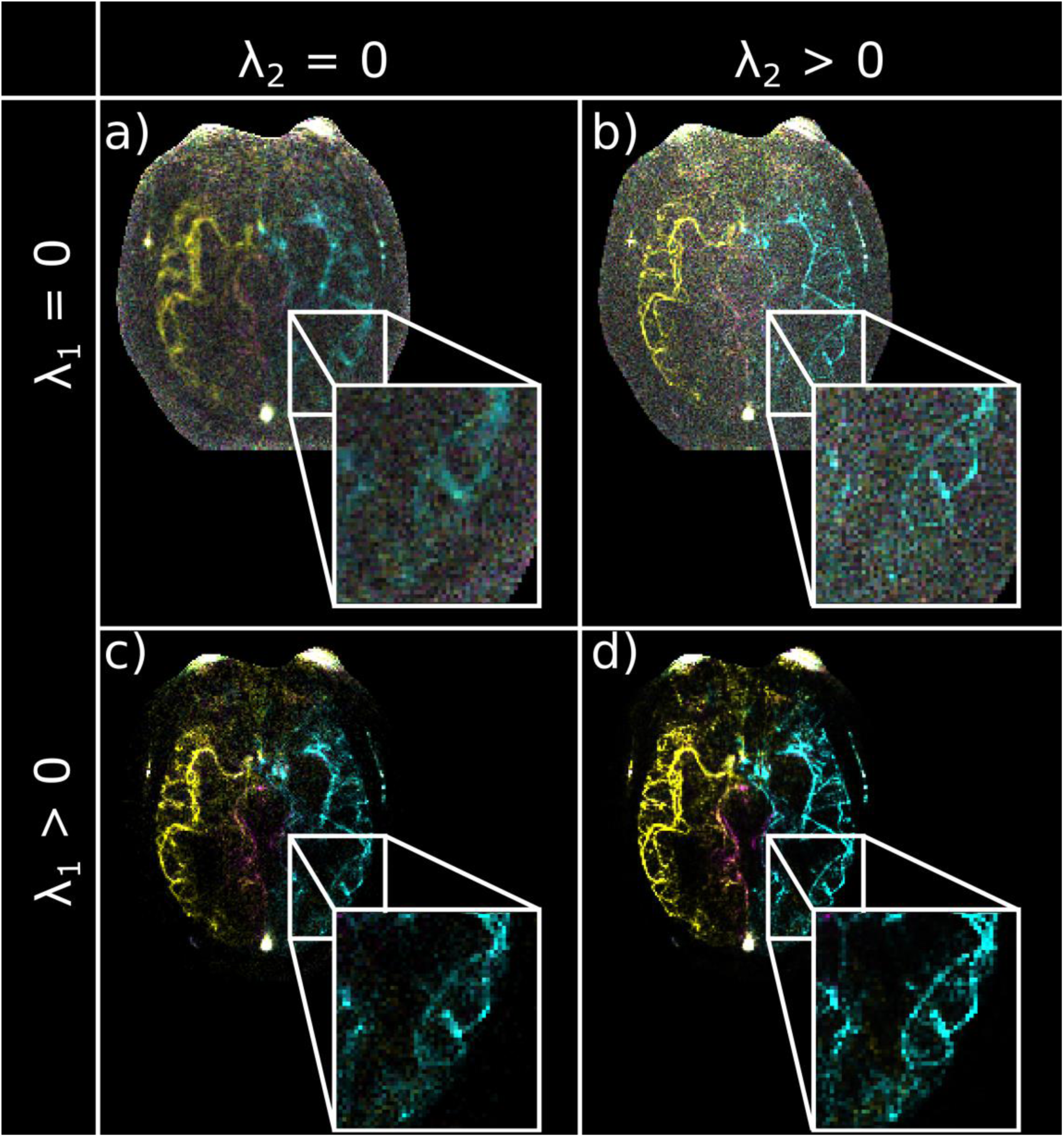
R = 34 VE-ASL reconstruction (temporal average) with a) no regularization, b) only L2 temporal smoothness constraint, c) only L1 sparsity constraint, d) both regularizing terms included. The L2-term sharpens the vessels and the L1-term denoises the background. Both regularizing terms improve the overall reconstruction quality.

### 3.4. Comparison of VE and nonVE at matched scan time

Generally, no statistically significant difference in image quality was found between VE and scan time matched nonVE images despite VE requiring a factor of two higher undersampling. A single exception was the high acceleration RICA where the nonVE correlation coefficients were marginally higher (P < 0.01). The in vivo results are summarized in Figure 8.

**Figure 8.**
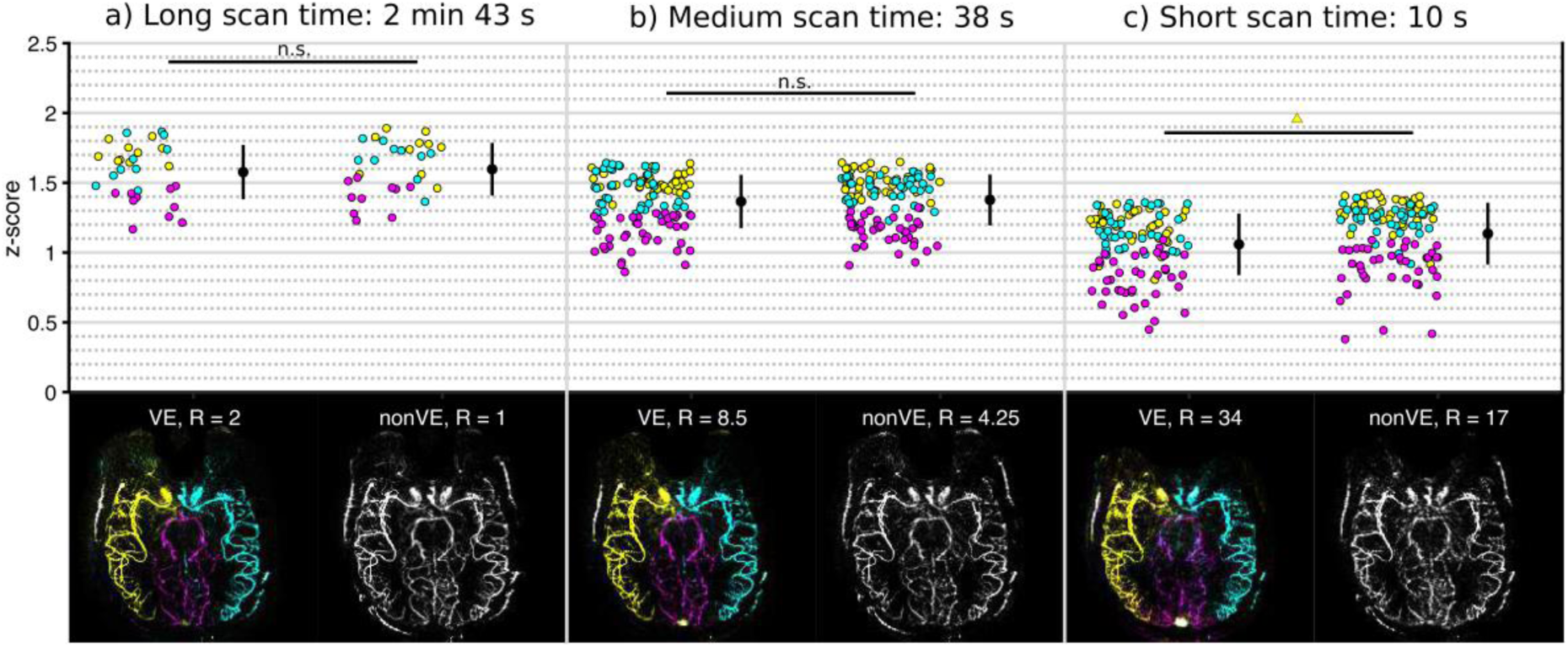
In-vivo reconstructions: a) fully sampled nonVE vs. R = 2 VE, b) moderate acceleration (R = 4.25 nonVE vs. R = 8.5 VE), c) high acceleration (R = 17 nonVE vs. R = 34 VE). The bottom row shows a single example of the reconstruction quality for one subsets.

Qualitatively the VE images also looked comparable with the nonVE images acquired at the same acquisition time (one example subject shown in Figure 8, reconstructions of the other subjects are available in Supporting Information Figures S2-6. At high R a loss of faint features compared with the ground truth was apparent for both nonVE and VE and some subsets included artifacts potentially caused by motion (Animations of the dynamic blood flow are available in the Supporting Information, Animations S7-11).

The temporal dynamics were also generally well conserved across acceleration factors. The signal was preserved in the late frames, but at the highest acceleration factors some residual aliasing was also present in the later frames (Figure 9) for both the nonVE and VE images. Figure 10 shows the temporal profile of the signal averaged in two 3×3 voxel regions in proximal and distal vessels in an example subject. In the distal vessel the SNR is lower and the temporal signal is noisier even in the ground truth case. The temporal regularization smooths the signal but preserves overall shape.

**Figure 9.**
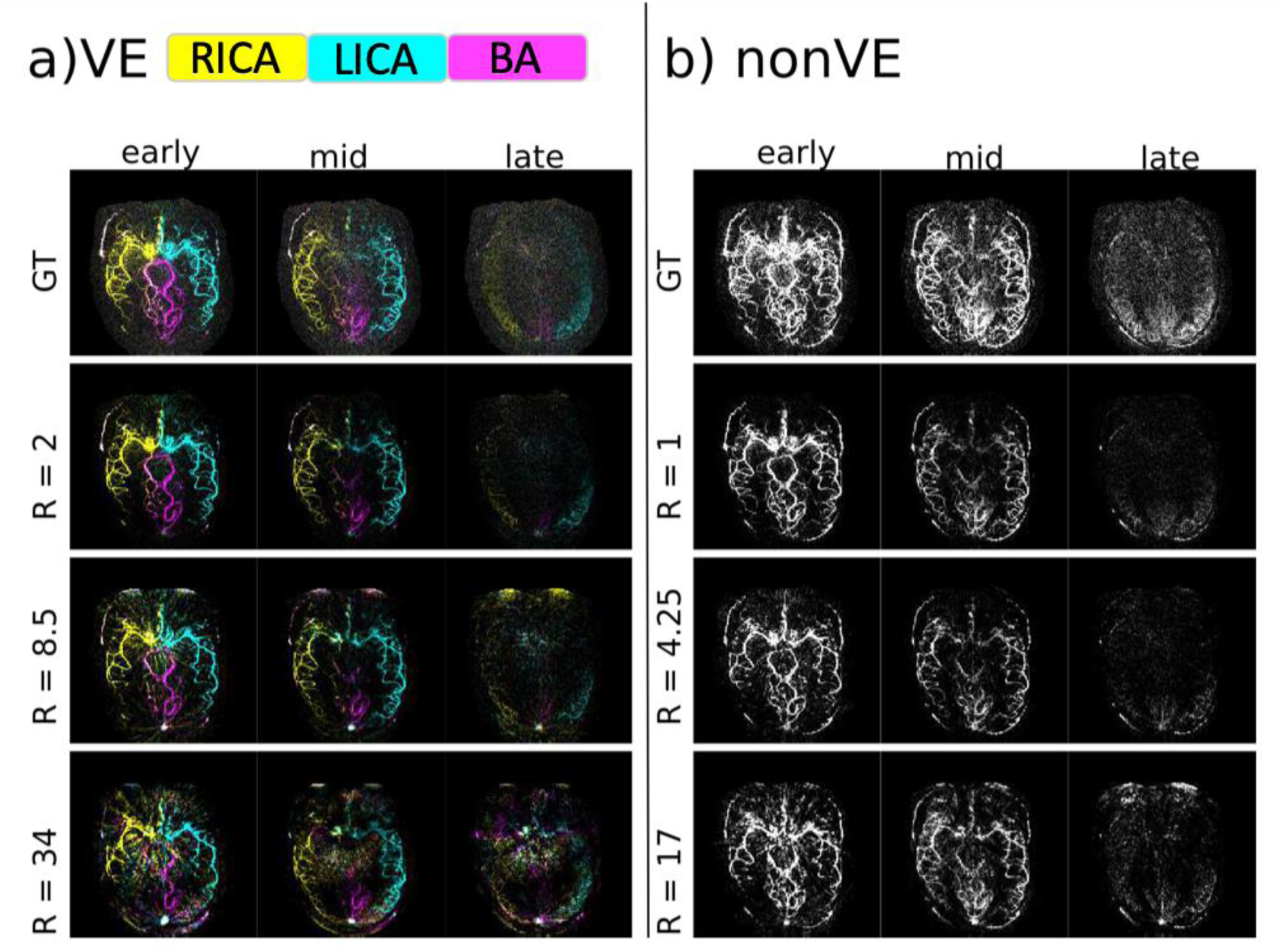
Temporal dynamics in an example subject at varying acceleration factors for a) a VE acquisition and b) a nonVE acquisition. The early time point is frame 1, the mid timepoint is frame 6 and the late time point is frame 12. Animations can be found in the Supporting Information (Animations S7-11).

**Figure 10.**
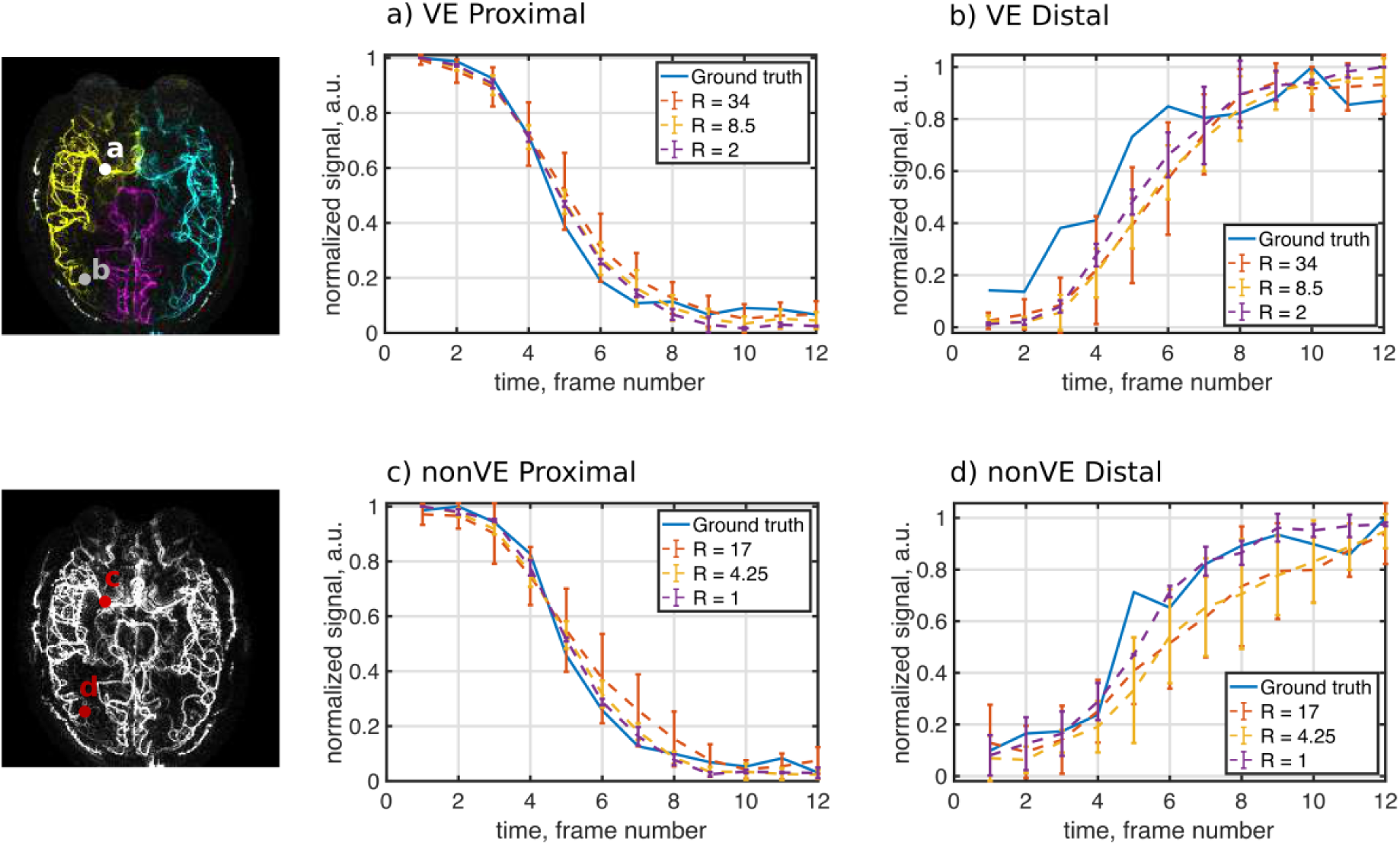
Temporal profile in two regions of interest, in a proximal vessel (a) and c)), and in a distal vessel (b) and d)) with blood supply from the RICA in one subject. The errorbars indicate the standard deviation of the signal measured from reconstructions of different subsets of the raw data at each acceleration factor.

## 4. Discussion and conclusions

In this study we have shown that the additional sparsity provided by vessel-encoding allows us to generate vessel-selective dynamic angiograms in the same time required for conventional ASL angiography without a loss of image quality.

### 4.1. The role of relative sparsity in CS and VE-ASL

As hypothesized, the simple simulation experiment showed that relative sparsity (proportion of non-zero voxels to total number of voxels reconstructed) alone, rather than absolute number of non-zero elements, can drive a L_1_-regularized reconstruction. This agrees with underlying theory of compressed sensing (15). How much relative sparsity drives the reconstruction quality compared to other factors such as SNR, and the spatial distribution of non-zero voxels, are topics worth further consideration. The theoretical and practical limitations of this method to get extra information “for free” in a system where relative sparsity increases should be further explored beyond this initial feasibility study.

One potential extension is applying this method to VE-ASL with labeling above the Circle of Willis, which would require more encodings because there are more vessel branches, but each decoded image would be sparser. This would theoretically allow for higher acceleration. However, practical issues such as with achieving an ideal Hadamard encoding for more complicated vessel geometries, might limit the improvements that the increased sparsity alone buys. A strategy for optimizing encodings for complex geometries has, however, been proposed previously (26).

To further aid reconstruction, the non-sparse nature of the static tissue needs to be considered. Although good images of the static tissue signal are not necessary from a clinical perspective, the static tissue component needs to be reconstructed well in order to correctly decode the blood signal. In this work the static tissue signal was treated just like the blood signals in reconstruction. It did not seem to hinder the reconstruction, as good results were achieved. However, further improvements could be achieved by applying a sparsifying transform such as the wavelet transform or spatial total variation constraints to it as it is not intrinsically sparse in image space like the blood vessels. In similar work on nonVE pulsed ASL (PASL) angiography wavelet regularization on the control image was reported to improve vessel delineation (27) By transforming the static tissue to a sparser domain, it should allow for better reconstruction in a compressed sensing framework.

### 4.2. Regularization in the temporal domain

In the same vein as increasing the spatial dimensionality by increasing the number of vessels encoded, the dynamic acquisition also allows you to take advantage of structure and redundancy in the temporal domain. In this work, the L_2_ norm of temporal finite differences was used to regularize the reconstruction. It was easily incorporated into the FISTA framework, and greatly improved the reconstruction results beyond the L_1_ spatial sparsity alone that mainly denoised the signal from both noise and noise-like aliasing. We observed that the introduction of the temporal regularization in particular improved spatial delineation, which could be explained by sharing information across frames that have different k-space sampling patterns, more precise estimation of the outer regions of k-space is possible. In previous works, temporal constraints have been enforced by using sliding-window acquisitions (28) or with compressed sensing using L_1_ constraints in temporal total variation (TV) frameworks (29), or in the temporal frequency domain (30). These approaches can, however, have unwanted effects such as increased blurring for the sliding-window method, temporal stair-casing artifacts for TV (31), or artificially introduced periodic behaviour in the temporal frequency domain. Whether these approaches could improve reconstruction for VE-ASL angiography could be studied further. Model based non-linear reconstruction is another option that has been explored in perfusion ASL (32). Using the model of temporal evolution of ASL angiography presented in (16), the reconstruction could be improved further and physiological metrics quantified.

### 4.3. Tuning the regularization factors

Among the five volunteers scanned in vivo the difference in optimal regularization factors was marginal and there was considerable overlap among their optimal regions. This suggests that once the reconstruction has been optimized for an acceleration factor and imaging protocol it is robust. However, because the optimal regularization factors depend on the properties of the data itself (inherent sparsity and temporal smoothness), one has to be careful with extending this conclusion to patients and other populations. There was some variability in head size, angle of imaging slab, and one of the volunteers exhibited considerable mixing of RICA and BA blood due to an asymmetrical Circle of Willis configuration, but for example patients with arteriovenous malformations could have considerably lower image domain sparsity due to the presence of additional abnormal vessels that can affect optimal regularization factors considerably. Further studies in appropriate patient groups are required to determine how generalizable this result is.

In this study the Pearson’s correlation coefficient, *r*, was used to objectively optimize the regularization parameters and define the quality of the reconstruction. We found the Pearson’s correlation coefficient to be a robust metric that corresponded well with visual perception of image quality, which was not the case for other metrics that were tested in preliminary work. Normalized-root-mean-square-error (NRMSE) is a straightforward metric but it does not correspond well with perceptual quality, as also reported in (33). Structural similarity index (SSIM) (34) that was developed specifically to correspond with visual perception worked well within a data set to e.g. tune the parameters, but was not suitable to compare different datasets as it was sensitive to the absolute scaling of the signal. The masking improved robustness as otherwise the result was mainly driven by large areas with no signal and regions containing artifacts such as eye motion. Tuning could also be done manually by experts to balance specificity (i.e. noise removal) and sensitivity (preservation of faint signals) for optimal clinical utility.

### 4.4. Feasibility of accelerating VE and nonVE ASL

The feasibility of acquiring vessel-encoded images without increasing scan time compared to fully sampled nonVE has here been clearly demonstrated in these results. Similar image quality was obtained with R = 2 VE and R = 1 nonVE imaging in vivo. Further reductions in scan time has also been shown to be possible, both for nonVE and VE ASL, with matched image quality. At the highest acceleration factor (R = 17 for nonVE and 34 for VE) the main features were still visible with scan times as short as 10 s. Although some loss of faint features and artifacts in the later frames with little to no signal was observed for high R, the required image quality will depend on the clinical application of the technique. For example, if the scan is acquired to add information about mixing of blood from different sources to other angiographic images a highly accelerated scan of lower quality might be sufficient, whereas if it is to be used diagnostically on its own, moderate acceleration factors might be more appropriate.

The small but statistically significant differences in correlation coefficient, *r*, between VE and nonVE at matched scan times that were measured both in simulations and in vivo were small compared with the effect of changing scan time or adding noise. Because the significant differences at high R varied in being in favor of VE and nonVE, and differences in qualitative assessment of the image quality small, this does not negate the overall conclusion of this study that VE and nonVE images of similar quality can be achieved at matched scan times.

For many clinical applications it would be desirable to extend this technique to 3D. The main reason 2D imaging of a single slab was chosen for this feasibility study, was to be able to acquire ground truth images in reasonable scan times. The extension to 3D is straightforward using a 3D golden angle radial sampling scheme (35) instead of the in-plane version. However, to fully sample VE ASL in 3D at the same resolution would take well over eight hours so no fully sampled ground truth could have been acquired for quantitative assessment. Extending to 3D will increase the relative sparsity further and should therefore allow for higher acceleration.

In conclusion, the lack of any meaningfully image quality differences between VE and nonVE data at matched scan times means that vessel-selective information can be acquired in this way with no cost of scan time or data quality, despite the higher undersampling factors required.

## References

1. Kaufmann TJ, Huston J, Mandrekar JN, Schleck CD, Thielen KR, Kallmes DF. Complications of Diagnostic Cerebral Angiography: Evaluation of 19 826 Consecutive Patients ^1^. Radiology 2007;243:812–819 doi: 10.1148/radiol.2433060536.

2. Grobner T. Gadolinium –a specific trigger for the development of nephrogenic fibrosing dermopathy and nephrogenic systemic fibrosis? Nephrol. Dial. Transplant. 2006;21:1104–1108 doi: 10.1093/ndt/gfk062.

3. Kanda T, Ishii K, Kawaguchi H, Kitajima K, Takenaka D. High Signal Intensity in the Dentate Nucleus and Globus Pallidus on Unenhanced T1-weighted MR Images: Relationship with Increasing Cumulative Dose of a Gadolinium-based Contrast Material. Radiology 2014;270:834–841 doi: 10.1148/radiol.13131669.

4. Tedeschi E, Caranci F, Giordano F, Angelini V, Cocozza S, Brunetti A. Gadolinium retention in the body: what we know and what we can do. Radiol. Med. (Torino) 2017;122:589–600 doi: 10.1007/s11547-017-0757-3.

5. Kanal E. Gadolinium based contrast agents (GBCA): Safety overview after 3 decades of clinical experience. Magn. Reson. Imaging 2016;34:1341–1345 doi: 10.1016/j.mri.2016.08.017.

6. Okell TW, Schmitt P, Bi X, et al. Optimization of 4D vessel-selective arterial spin labeling angiography using balanced steady-state free precession and vessel-encoding. NMR Biomed. 2016;29:776–786 doi: 10.1002/nbm.3515.

7. Bang OY, Goyal M, Liebeskind DS. Collateral Circulation in Ischemic Stroke: Assessment Tools and Therapeutic Strategies. Stroke 2015;46:3302–3309 doi: 10.1161/STROKEAHA.115.010508.

8. Warmuth C, Ruping M, Forschler A, et al. Dynamic Spin Labeling Angiography in Extracranial Carotid Artery Stenosis. Am. J. Neuroradiol. 2005;26:1035–1043.

9. Wong EC. Vessel-encoded arterial spin-labeling using pseudocontinuous tagging. Magn. Reson. Med. 2007;58:1086–1091 doi: 10.1002/mrm.21293.

10. Dai W, Garcia D, de Bazelaire C, Alsop DC. Continuous Flow Driven Inversion for Arterial Spin Labeling Using Pulsed Radiofrequency and Gradient Fields. Magn. Reson. Med. 2008;60:1488–1497 doi: 10.1002/mrm.21790.

11. Helle M, Norris DG, Rüfer S, Alfke K, Jansen O, van Osch MJP. Superselective pseudocontinuous arterial spin labeling. Magn. Reson. Med. 2010;64:777–786 doi: 10.1002/mrm.22451.

12. Dai W, Robson PM, Shankaranarayanan A, Alsop DC. Modified pulsed continuous arterial spin labeling for labeling of a single artery. Magn. Reson. Med. 2010;64:975–982 doi: 10.1002/mrm.22363.

13. Lustig M, Donoho D, Pauly JM. Sparse MRI: The application of compressed sensing for rapid MR imaging. Magn. Reson. Med. 2007;58:1182–1195 doi: 10.1002/mrm.21391.

14. Donoho DL. Compressed sensing. IEEE Trans. Inf. Theory 2006;52:1289–1306 doi: 10.1109/TIT.2006.871582.

15. Candes EJ, Romberg J, Tao T. Robust uncertainty principles: exact signal reconstruction from highly incomplete frequency information. IEEE Trans. Inf. Theory 2006;52:489–509 doi: 10.1109/TIT.2005.862083.

16. Okell TW, Chappell MA, Schulz UG, Jezzard P. A kinetic model for vessel-encoded dynamic angiography with arterial spin labeling. Magn. Reson. Med. 2012;68:969–979 doi: 10.1002/mrm.23311.

17. Fessler JA, Sutton BP. Nonuniform fast Fourier transforms using min-max interpolation. IEEE Trans. Signal Process. 2003;51:560–574 doi: 10.1109/TSP.2002.807005.

18. Fessler JA. Michigan Image Reconstruction Toolbox. Available at https://web.eecs.umich.edu/~fessler/code/ Accessed on 2018-02-26

19. Winkelmann S, Schaeffter T, Koehler T, Eggers H, Doessel O. An Optimal Radial Profile Order Based on the Golden Ratio for Time-Resolved MRI. IEEE Trans. Med. Imaging 2007;26:68–76 doi: 10.1109/TMI.2006.885337.

20. Okell TW. Combined angiography and perfusion using radial imaging and arterial spin labeling. Magn. Reson. Med. 2018 doi: 10.1002/mrm.27366.

21. Wu H, Block WF, Turski PA, Mistretta CA, Johnson KM. Noncontrast-enhanced three-dimensional (3D) intracranial MR angiography using pseudocontinuous arterial spin labeling and accelerated 3D radial acquisition. Magn. Reson. Med. 2013;69:708–715 doi: 10.1002/mrm.24298.

22. Buehrer M, Pruessmann KP, Boesiger P, Kozerke S. Array compression for MRI with large coil arrays. Magn. Reson. Med. 2007;57:1131–1139 doi: 10.1002/mrm.21237.

23. Walsh DO, Gmitro AF, Marcellin MW. Adaptive reconstruction of phased array MR imagery. Magn. Reson. Med. 2000;43:682–690 doi: 10.1002/(SICI)1522-2594(200005)43:5<682::AID-MRM10>3.0.CO;2-G.

24. Beck A, Teboulle M. A Fast Iterative Shrinkage-Thresholding Algorithm for Linear Inverse Problems. SIAM J. Imaging Sci. 2009;2:183–202 doi: 10.1137/080716542.

25. Fessler JA, Sangwoo Lee, Olafsson VT, Shi HR, Noll DC. Toeplitz-based iterative image reconstruction for MRI with correction for magnetic field inhomogeneity. IEEE Trans. Signal Process. 2005;53:3393–3402 doi: 10.1109/TSP.2005.853152.

26. Berry ESK, Jezzard P, Okell TW. An Optimized Encoding Scheme for Planning Vessel-Encoded Pseudocontinuous Arterial Spin Labeling: Optimized Encoding Scheme for VEASL. Magn. Reson. Med. 2015;74:1248–1256 doi: 10.1002/mrm.25508.

27. Zhou Z, Han F, Yu S, et al. Accelerated noncontrast-enhanced 4-dimensional intracranial MR angiography using golden-angle stack-of-stars trajectory and compressed sensing with magnitude subtraction. Magn. Reson. Med. 2018;79:867–878 doi: 10.1002/mrm.26747.

28. Barger AV, Block WF, Toropov Y, Grist TM, Mistretta CA. Time-resolved contrast-enhanced imaging with isotropic resolution and broad coverage using an undersampled 3D projection trajectory. Magn. Reson. Med. 2002;48:297–305 doi: 10.1002/mrm.10212.

29. Feng L, Grimm R, Block KT, et al. Golden-angle radial sparse parallel MRI: Combination of compressed sensing, parallel imaging, and golden-angle radial sampling for fast and flexible dynamic volumetric MRI: iGRASP: Iterative Golden-Angle RAdial Sparse Parallel MRI. Magn. Reson. Med. 2014;72:707–717 doi: 10.1002/mrm.24980.

30. Lustig M, Santos JM, Donoho DL. K-t SPARSE: High Frame Rate Dynamic MRI Exploiting Spatio-Temporal Sparsity. In: Proceedings of the 14th Annual Meeting of ISMRM.; 2006.

31. Jian Zhang, Shaohui Liu, Ruiqin Xiong, Siwei Ma, Debin Zhao. Improved total variation based image compressive sensing recovery by nonlocal regularization. In: 2013 IEEE International Symposium on Circuits and Systems (ISCAS2013). Beijing: IEEE; 2013. pp. 2836–2839. doi: 10.1109/ISCAS.2013.6572469.

32. Zhao L, Fielden SW, Feng X, Wintermark M, Mugler JP, Meyer CH. Rapid 3D dynamic arterial spin labeling with a sparse model-based image reconstruction. NeuroImage 2015;121:205–216 doi: 10.1016/j.neuroimage.2015.07.018.

33. Akasaka T, Fujimoto K, Yamamoto T, et al. Optimization of Regularization Parameters in Compressed Sensing of Magnetic Resonance Angiography: Can Statistical Image Metrics Mimic Radiologists’ Perception? PLOS ONE 2016;11:e0146548.doi: 10.1371/journal.pone.0146548.

34. Wang Z, Bovik AC, Sheikh HR, Simoncelli EP. Image Quality Assessment: From Error Visibility to Structural Similarity. IEEE Trans. Image Process. 2004;13:600–612 doi: 10.1109/TIP.2003.819861.

35. Chan RW, Ramsay EA, Cunningham CH, Plewes DB. Temporal stability of adaptive 3D radial MRI using multidimensional golden means. Magn. Reson. Med. 2009;61:354–363 doi: 10.1002/mrm.21837.

