## Supporting Information for "Highly Accelerated Vessel-Selective Arterial Spin Labelling Angiography using Sparsity and Smoothness Constraints"

*Supporting Information Table S1* – Optimal regularisation factors for a) in vivo and b) simulation

| **a) IN VIVO** | Subj 1 | | Subj 2 | | Subj 3 | | Subj 4 | | Subj 5 | | Overall | |
|---|---|---|---|---|---|---|---|---|---|---|---|---|
| | $\lambda_1$ | $\lambda_2$ | $\lambda_1$ | $\lambda_2$ | $\lambda_1$ | $\lambda_2$ | $\lambda_1$ | $\lambda_2$ | $\lambda_1$ | $\lambda_2$ | $\lambda_1$ | $\lambda_2$ |
| nonVE R = 1 | 4.00E-06 | 4 | 4.00E-06 | 4 | 4.00E-06 | 2 | 4.00E-06 | 2 | 5.00E-06 | 2 | 4.00E-06 | 2 |
| VE R = 2 | 7.00E-06 | 6 | 7.00E-06 | 6 | 7.00E-06 | 6 | 7.00E-06 | 6 | 7.00E-06 | 4 | 7.00E-06 | 6 |
| nonVE R = 4.25 | 4.00E-06 | 4 | 4.00E-06 | 4 | 5.00E-06 | 4 | 4.00E-04 | 3 | 6.00E-06 | 3 | 5.00E-06 | 3 |
| VE R = 8.5 | 6.00E-06 | 5 | 6.00E-06 | 4 | 6.00E-06 | 4 | 6.00E-06 | 5 | 7.00E-06 | 4 | 6.00E-06 | 4 |
| nonVE R = 17 | 3.00E-06 | 1.8 | 3.00E-06 | 2.2 | 4.00E-06 | 2.2 | 3.00E-06 | 1.6 | 4.00E-06 | 1.8 | 3.00E-06 | 1.8 |
| VE R = 34 | 1.00E-06 | 0.8 | 4.00E-06 | 2 | 4.00E-06 | 2 | 4.00E-06 | 2 | 5.00E-06 | 2 | 4.00E-06 | 1.8 |

| **b) SIMULATION** | $SNR_k = \infty$ | | $SNR_k$ = 185.7 | | $SNR_k$ = 92.8 | |
|---|---|---|---|---|---|---|
| | $\lambda_1$ | $\lambda_2$ | $\lambda_1$ | $\lambda_2$ | $\lambda_1$ | $\lambda_2$ |
| nonVE R = 1 | 0 | 0 | 1.00E-06 | 2 | 4.00E-06 | 2 |
| VE R = 2 | 0 | 0 | 1.00E-06 | 2 | 7.00E-06 | 4 |
| nonVE R = 4.25 | 1.00E-06 | 0 | 3.00E-06 | 1 | 5.00E-06 | 3 |
| VE R = 8.5 | 1.00E-06 | 0 | 3.00E-06 | 2 | 7.00E-06 | 3 |
| nonVE R = 17 | 1.00E-06 | 0.2 | 2.00E-06 | 0.6 | 4.00E-06 | 1.4 |
| VE R = 34 | 1.00E-06 | 0.2 | 2.00E-06 | 0.6 | 5.00E-06 | 1.4 |

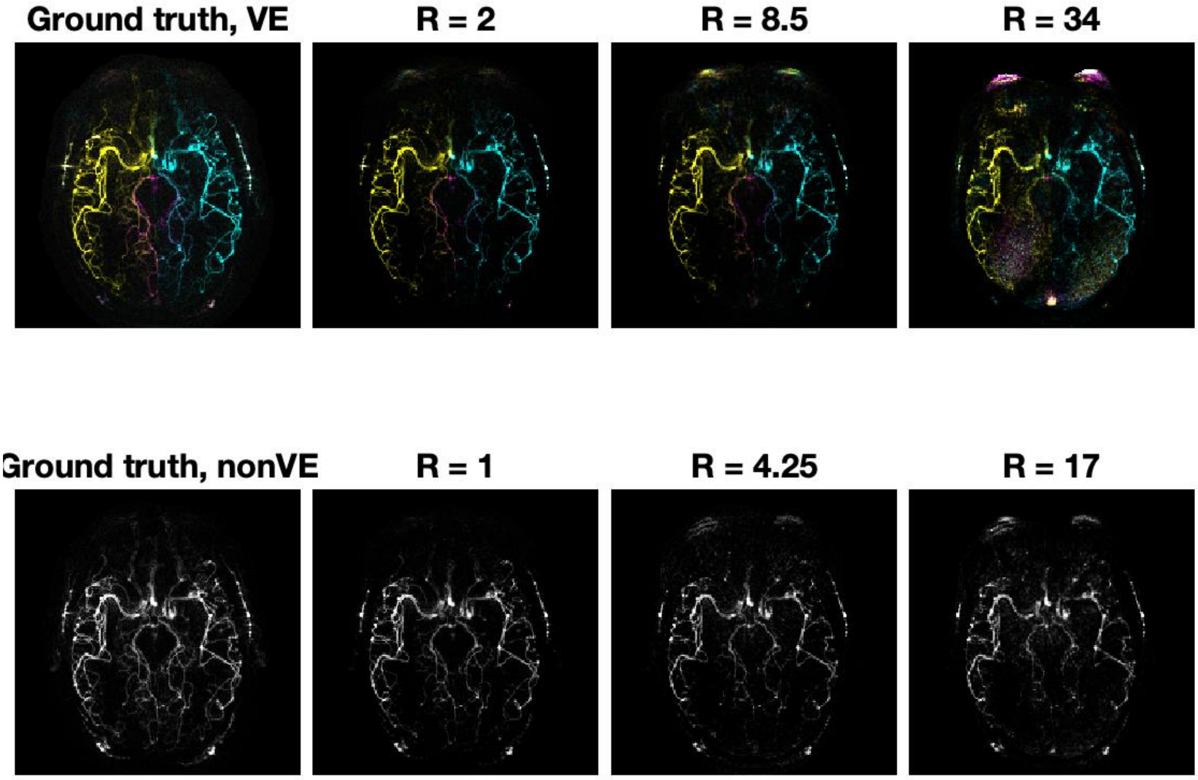


*Figure S2* – Time averaged in-vivo reconstruction of Subject 1: The top row shows the VE reconstruction with blood originating in the RICA in yellow, The LICA in cyan, and BA in magenta at varying acceleration factors. The bottom row shows the time matched nonVE images.

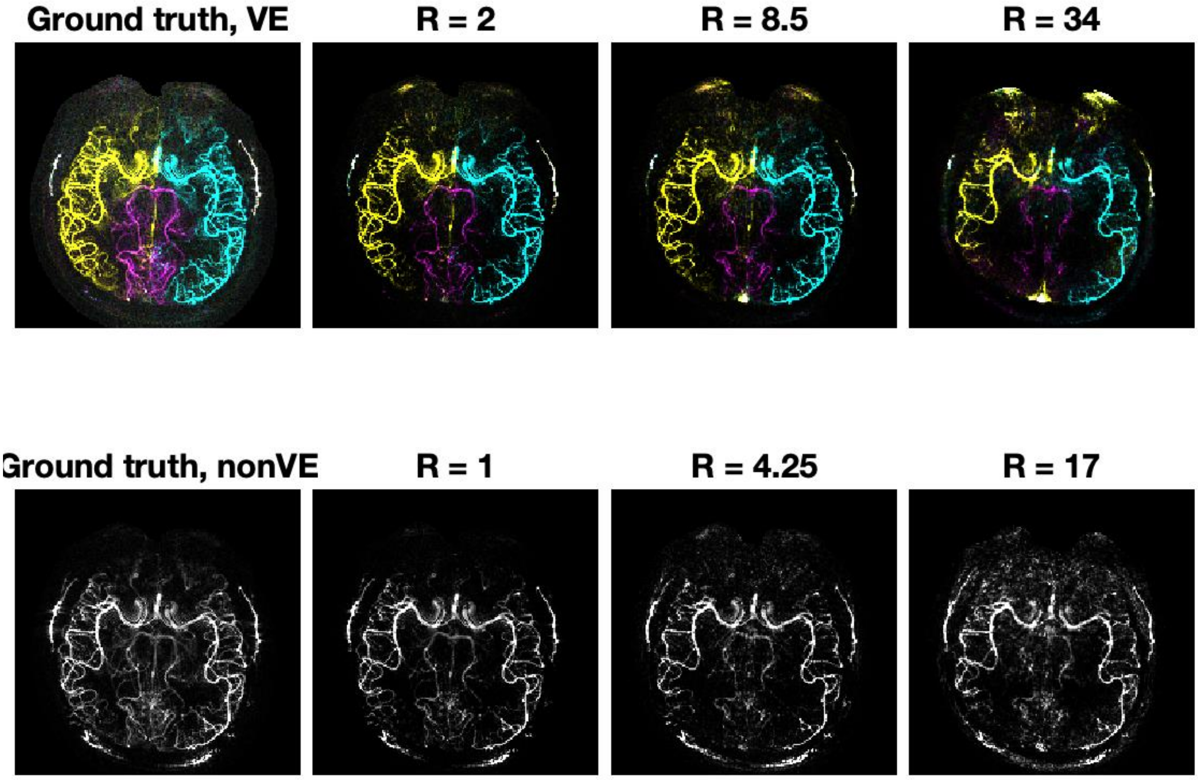


*Figure S3 -* Time averaged in-vivo reconstruction of Subject 2: The top row shows the VE reconstruction with blood originating in the RICA in yellow, The LICA in cyan, and BA in magenta at varying acceleration factors. The bottom row shows the time matched nonVE images.

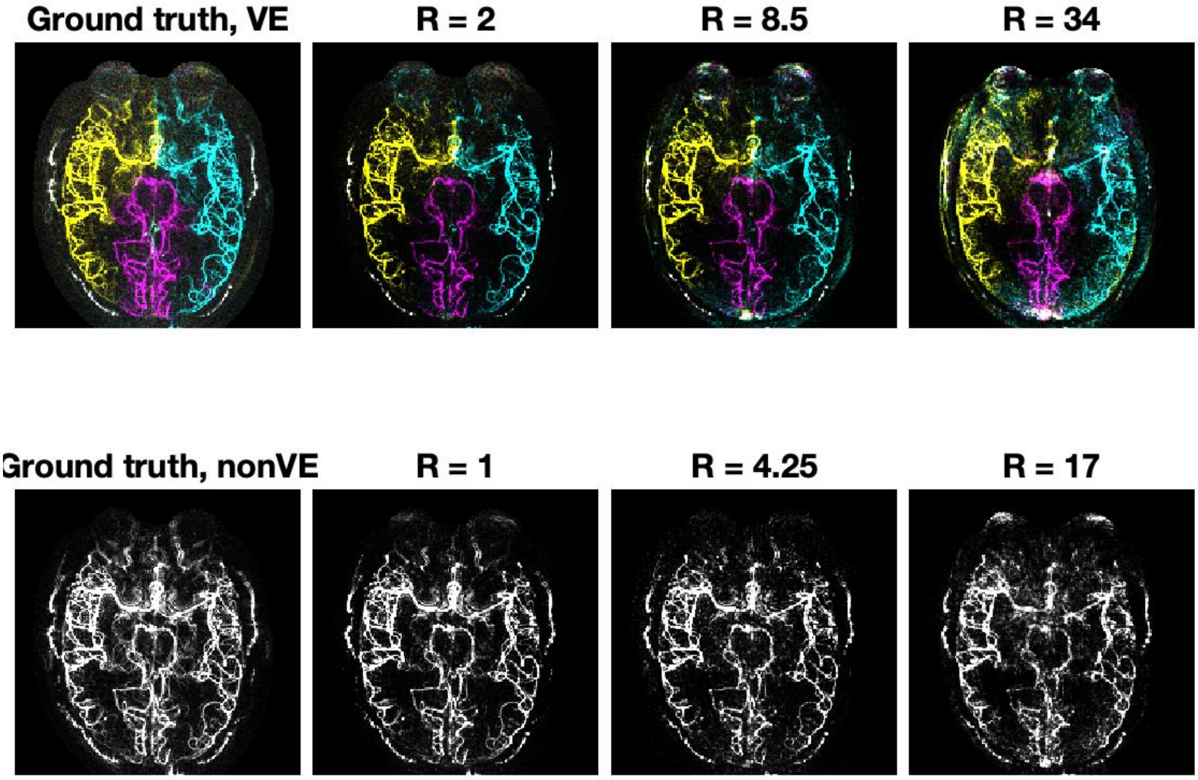


*Figure S4* - Time averaged in-vivo reconstruction of Subject 3: The top row shows the VE reconstruction with blood originating in the RICA in yellow, The LICA in cyan, and BA in magenta at varying acceleration factors. The bottom row shows the time matched nonVE images.

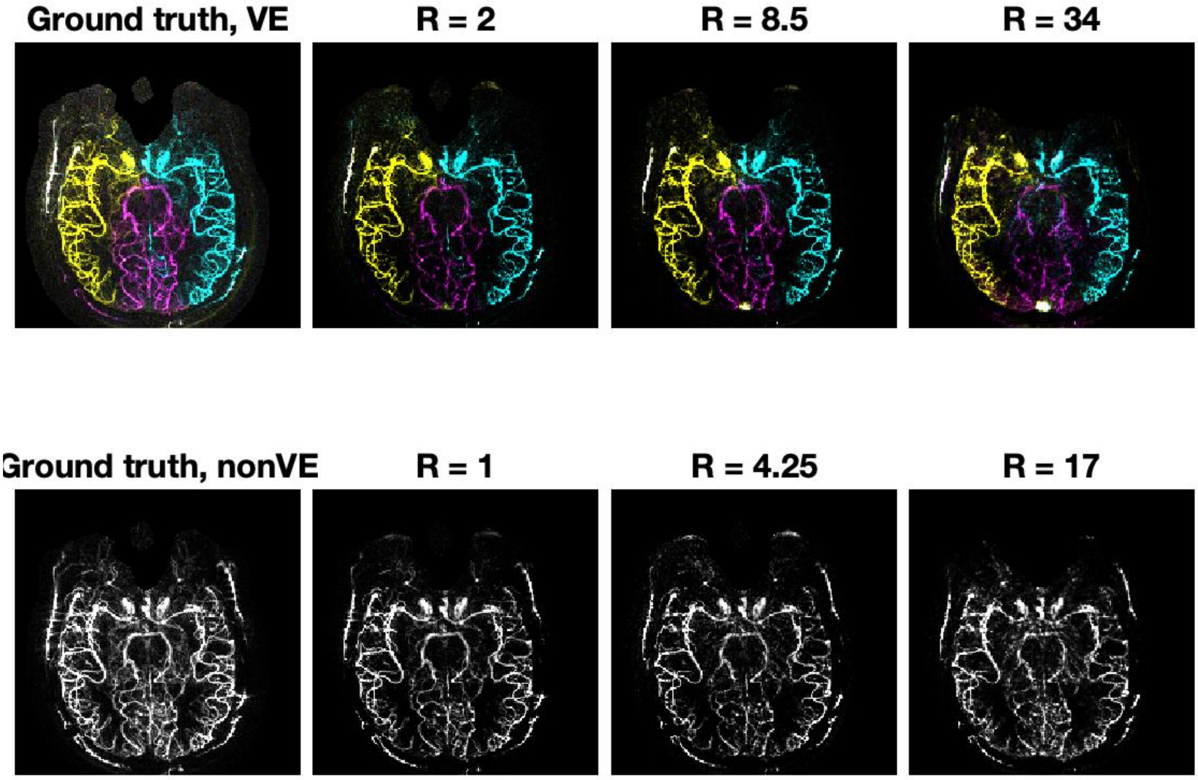


*Figure S5* - Time averaged in-vivo reconstruction of Subject 4: The top row shows the VE reconstruction with blood originating in the RICA in yellow, The LICA in cyan, and BA in magenta at varying acceleration factors. The bottom row shows the time matched nonVE images.

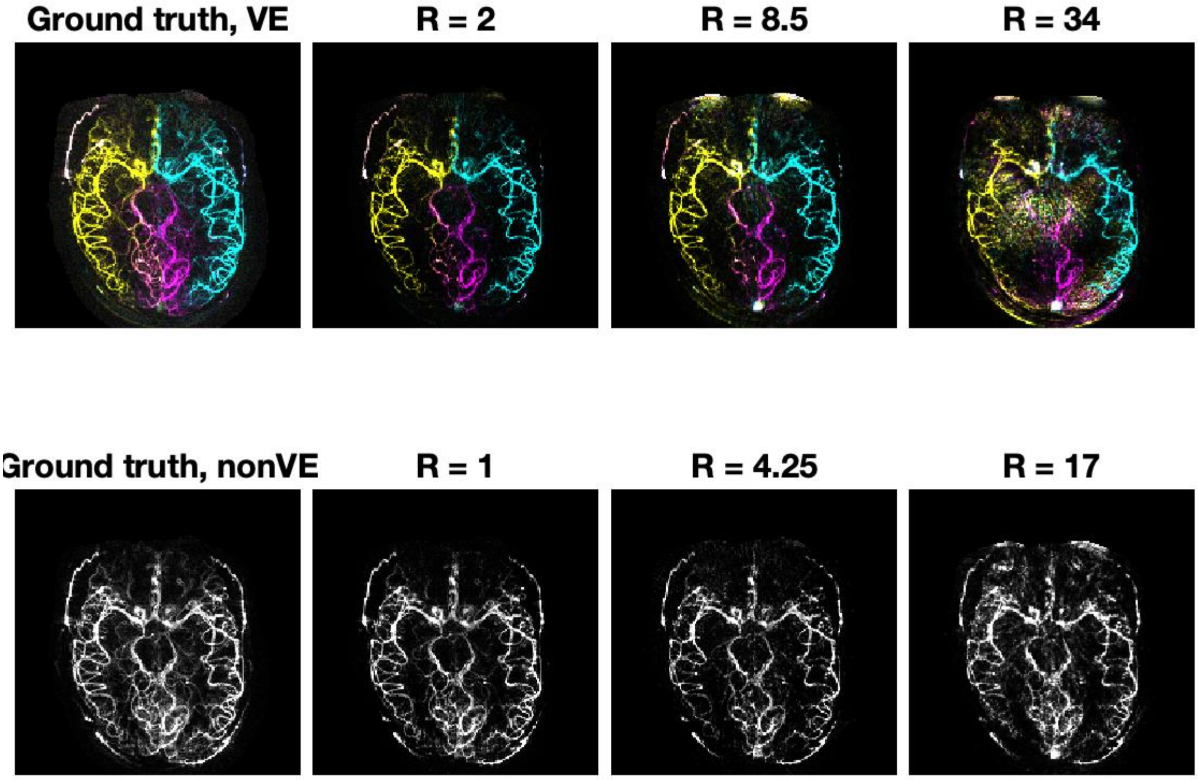


*Figure S6* – Time averaged in-vivo reconstruction of Subject 5: The top row shows the VE reconstruction with blood originating in the RICA in yellow, The LICA in cyan, and BA in magenta at varying acceleration factors. The bottom row shows the time matched nonVE images.
